# From proteins to nanoparticles: domain-agnostic predictions of nanoscale interactions

**DOI:** 10.1101/2022.08.09.503361

**Authors:** Jacob Saldinger, Matt Raymond, Paolo Elvati, Angela Violi

**Affiliations:** Chemical Engineering, University of Michigan, Street, Ann Arbor, 48109-2125, Michigan, USA; Electrical Engineering and Computer Science, University of Michigan, Street, Ann Arbor, 48109-2125, Michigan, USA; Mechanical Engineering, University of Michigan, Street, Ann Arbor, 48109-2125, Michigan, USA

**Keywords:** Neural Networks, Proteins, Nanoparticles, Dimensionality Reduction, Coarse-Graining

## Abstract

The accurate and rapid prediction of generic nanoscale interactions is a challenging problem with broad applications. Much of biology functions at the nanoscale, and our ability to manipulate materials and engage biological machinery in a purposeful manner requires knowledge of nano-bio interfaces. While several protein-protein interaction models are available, they leverage protein-specific information, limiting their abstraction to other structures. Here, we present NeCLAS, a general, and rapid machine learning pipeline that predicts the location of nanoscale interactions, providing human-intelligible predictions. Two key aspects distinguish NeCLAS: coarsegrained representations, and the use of environmental features to encode the chemical neighborhood. We showcase NeCLAS with challenges for protein-protein, protein-nanoparticle and nanoparticle-nanoparticle systems, demonstrating that NeCLAS replicates computationally- and experimentally-observed interactions. NeCLAS outperforms current nanoscale prediction models and it shows cross-domain validity. We anticipate that our framework will contribute to both basic research and rapid prototyping and design of diverse nanostructures in nanobiotechnology.

---

Many technological, biological, and natural phenomena are governed by molecular and nanoscale interactions and the processes that occur at their interfaces [1–4]. Protein-protein interactions (PPIs) play a crucial role in cellular functions and biological processes in all organisms, from mediating selectivity along signaling pathways to understanding of infection mechanisms, to development of treatments and therapies. Similarly, protein-nanoparticle interactions provide knowledge about the bio-reactivity of nanoparticles and their applications in nanodiagnostics, nanotherapy, and nanomedicine. However, to tailor these interactions, a comprehensive knowledge of how nanomaterials interact with biological systems is critical.

In recent years, data-driven machine learning (ML) methods have emerged as powerful tools to provide insight into the mechanisms and chemistry of nanoscale interactions, overcoming the cost and complexity of experiments [5–7] and simulations [8–12], without the need of *a priori* knowledge of physics- and template-based methods [13–15].

Partner-independent ML methods predict interaction sites for a target structure, regardless of the complementary nanostructure, and can successfully predict protein-ligand [16] and protein-protein [17–19] interactions. These methods identify features that correlate with the tendency of the target protein to interact with arbitrary nanostructures, but do not consider the properties of the second molecule (partner) directly. This approach is data-efficient, but pairwise information from interacting partners is often highly relevant for specific molecules and results in improved predictions [20, 21]. To address this limitation, partner-specific methods were developed to predict whether a subunit (*e.g*., protein residue) of one structure interacts with a specific subunit of another complex [20, 22, 23]. Crucially, by using curated datasets [20, 24] that include diverse structures and account for homology, partner-specific methods were shown to successfully predict the local pairwise residue interactions that control global protein-protein aggregation.

Despite this progress, most of the current approaches are specifically designed for proteins and are not immediately generalizable. As these methods use properties of the individual amino acids or rely on protein-specific characteristics (*i.e*., properties derived from sequence and residue conservation), they cannot be straightforwardly extended to molecules that lack these motifs, even when they share other physical and chemical features [1, 25–27]. Similarly, current ML methods for predicting nanoparticle-protein interactions use application-specific properties and are limited by small training datasets [28–30], which limits the cross-domain validity of the resulting ML models, and requires a new model for every application.

To relax this specificity, we introduce NeCLAS, **Ne**ural **C**oarse-graining with **L**ocation **A**gnostic **S**ets, a flexible and generalized machine learning approach for predicting partner-specific nanoscale interactions. NeCLAS has two main features. The first is a generalized, atomistically-derived coarse-graining method to generate a rototranslational equivariant representation of nanoparticles and macromolecules. The second is a permutation invariant deep neural network that predicts pairwise interactions between the coarse-grained sites of two different molecules. We showcase NeCLAS with 3 increasingly complex prediction challenges: (1) binding site for protein-nanoparticle interactions; (2) dynamic characteristics of nanoparticle-protein interactions, and (3) nanoparticle-nanoparticle interactions and their tendency to self-assemble.

Without any protein-specific descriptors, testing on curated datasets shows that NeCLAS outperforms state-of-the-art protein-nanoparticle prediction methods, is competitive with the best protein-specific methods, and shows potential in predicting nanoparticle-nanoparticle interactions. Overall, NeCLAS demonstrates a versatile framework for interaction predictions across multiple domains with a reduced computational footprint. Our conceptual framework finds applications in various fields, from biologists who search for interactions between proteins, to materials scientists who can design and engineer nanoparticles for targeted applications, to the broad range of nanobiotechnology.

## Results

### NeCLAS: a domain-agnostic pipeline

ML model development can often be abstracted into two main steps: creating a learnable representation of real-world data, and using this representation to train a predictive model (Fig. 1a). In NeCLAS, the first step is accomplished by converting atomistic information to lower-dimensional coarse-grained (CG) structures and then computing properties for each CG site, accounting for both local characteristics and chemical neighborhood. The second step is accomplished by training a permutation invariant deep neural network to predict pairwise interactions of these CG sites.

**Fig. 1.**
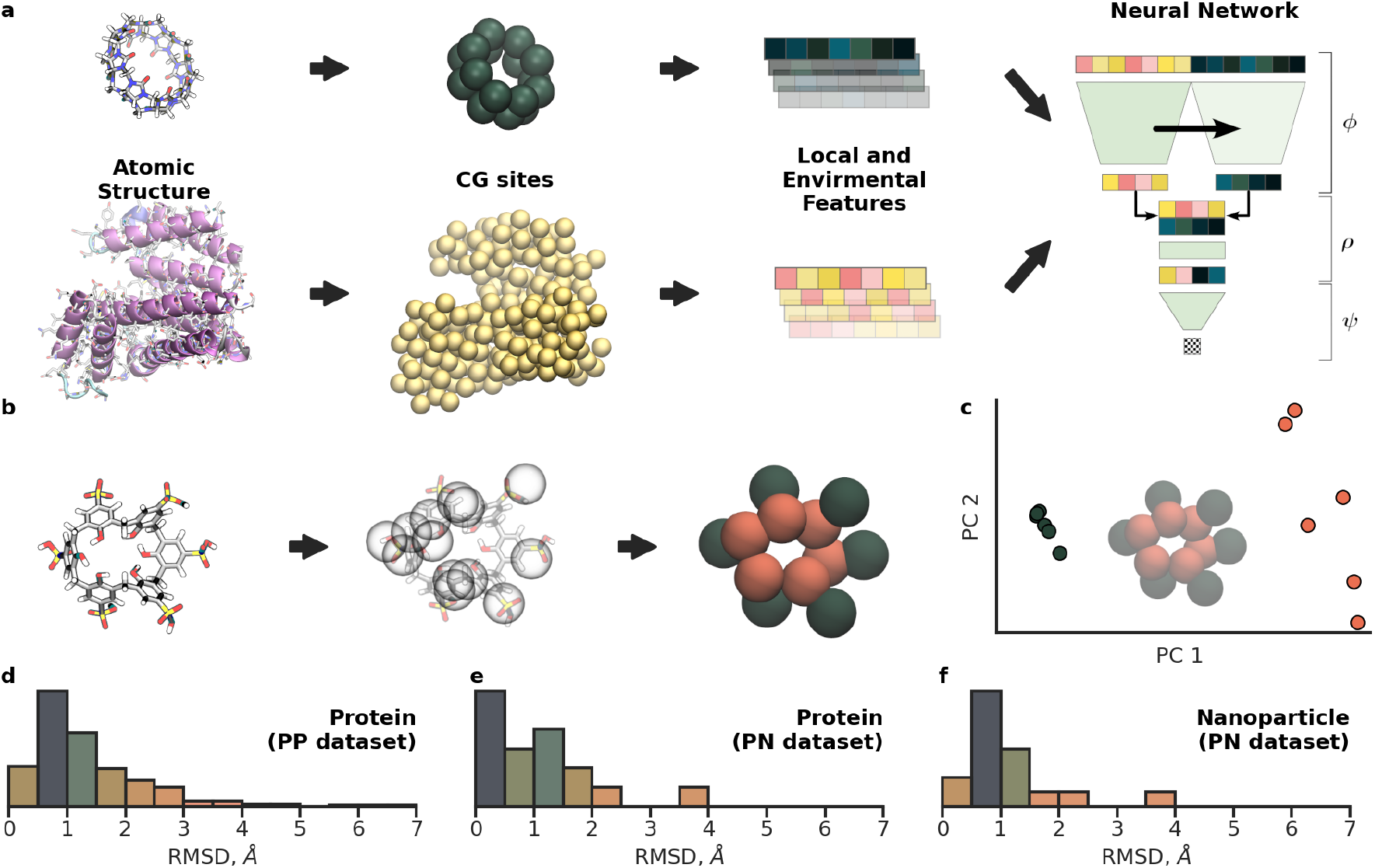
Methods and data. **a**, NeCLAS schematic. Reduced dimensionality representation (CG sites) and properties are derived from atomic structures (*e.g*., nanoparticles, top row, and proteins, bottom row); a set of the combined local (80) and environmental (400) features is then generated for each pairwise interactions and used for training and testing the Neural Network. **b**, Schematic of the coarse-graining procedure applied on *p*-sulfocalix[6]arene (*p*-Sclx_6_). CG subunits centers are randomly placed within the coordinate space of a starting molecule, then iteratively shifted to match a target property spatial distribution. **c**, First two principal components of the feature set obtained for *p*-Sclx_6_. Colors correspond to CG sites shown on the CG molecule in left panel of **c. d**, Distribution of RMSD between unbound and bound proteins in the protein-protein (PP) dataset (DBD version 5). One structure (PDB: 1IRA) was omitted for clarity (RMSD = 8.36). **e-f**, Distribution of RMSD between unbound and bound proteins and nanoparticles for the protein-nanoparticle (PN) dataset.

To obtain the CG sites (Fig. 1b), a predetermined number of sites is randomly initialized and iteratively optimized to match the atomic distribution of a given molecular property (*e.g*., mass) [31]. The resulting representation can be easily tailored to capture structural symmetries, especially when interpretability is a primary concern. For example, *p*-Sclx_6_ (Fig. 1b) is a para-sulfonate calixarene, composed of six repeating units with a positively-changed outer region and negatively-charged inner region [32]. By using 12 CG sites, the procedure consistently allocates two sets of sites that match the symmetry of the molecule. Indeed, *p*-Sclx_6_ has a hydrophobic core and anionic rim that facilitates protein recognition via entrapment of arginine or lysine side chains. These CG sites capture the underlying molecular properties, as shown by the two distinct clusters of the CG sites in the principal component analysis (Fig. 1c) of their local chemical features. While useful for interpretability, the specific choice of number of sites has a minimal impact on the accuracy of NeCLAS, as long as the extreme choices (*e.g*., one site per nanoparticle) are avoided (Supplementary Information, section 1).

Once the lower dimensionality representation is obtained, local and environmental features are computed for each site. We selected 80 CG-site properties (local) that are generally important for chemical and biological interactions, such as charge, shape, size, hydrogen bonding, depth, and surface exposure [33–38]. To capture the effect of the surrounding atoms, which are known to play an important role [20, 22, 39], we considered 400 properties weighted by spatial functions that provide a description of the molecular environment that is (globally) equivariant under translations and rotations [40, 41].

To evaluate our model, we specifically tailored our data and workflow to avoid common causes of artificially inflated estimates of the model performance. First, to ensure model generalization, NeCLAS utilizes a neural network that is invariant to the ordering of input sites by design. This structure provides a more stable prediction compared to permutation variant methods (Supplementary Information, section 4). As per standard ML practice, we kept a strict separation between train, validation (used to halt the neural network learning process), and test sets (used to evaluate model performance). We chose our datasets to avoid data redundancy, and assessed the performance of our model with increasingly stricter criteria to test the possibility of information leakage from the training set to the validation or test sets. Lastly, since proteins and nanoparticles may change conformations as they interact, we trained NeCLAS only on unbound structures, as the ultimate goal is to predict interactions for species with an unknown bound conformation [42]. For our datasets, these structural changes were quantified as the root mean squared deviations (RMSD) of the atomic positions of bound and unbound species. The distributions (Fig. 1d-f), show that during binding, conformational changes can be significant.

Following these guiding principles, for proteinprotein interactions, we choose the Docking Benchmark Dataset (DBD) version 5, a curated set of 230 experimental structures of nonredundant protein complexes in both bound and unbound form [24]. However, for protein-nanoparticles interactions no such dataset exists, and therefore we created a dataset from an existing set of data provided by Costanzo’ *et al*. [43], which contains organic nanoparticles (*e.g*., fullerenes, macrocages, calixarenes, cyclodextrins, cucurbiturils, and molecular tweezers). From this data, we generated both bound and unbound structures. Since this dataset is relatively small and structural redundancy cannot be avoided, we used it only for testing, preventing information leakage from similar substructures.

### Performance

Below we assess the performance of (NeCLAS) to predict nanoparticle-protein interactions in comparison with five different techniques: the very recently published generalized method (Unified [44]), and four binding residue prediction methods that are not partner specific, namely SPPIDER [39] (designed for protein-protein interactions), P2Rank [17] and COACH [45] (for protein-ligand binding), and Fpocket [46] (which identifies pockets in the protein geometry). The last four methods were not originally designed to predict protein-nanoparticle interactions, but they can be used as they are not partner-specific.

To evaluate each technique, we generated the receiver operating characteristic curve, which computes the fraction of true-positive to falsepositive interaction predictions at various discrimination thresholds. The area under this curve (AUC) quantifies the quality of a binary model’s predictions, and it is a commonly used evaluation metric in similar problems [18, 20, 22, 23]. In the typical formulation, AUC is computed using the entire testing dataset (AUC_all_); however, because nanoscale complexes can have different sizes, here we also use the median of values computed for each individual complex, AUC_comp_. AUC_comp_ reweights pairwise interactions to ensure equal contribution of each complex regardless of its size, which produces a more realistic metric for model performance on a new species. As NeCLAS and Unified do not predict interaction interfaces directly, we converted pairwise predictions to interface predictions using a scoring function [20], which considers interface membership using all the possible interactions with different weights 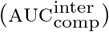.

Finally, to test the importance of coarsegrained pairwise information on the outcome of predictions, we also included a non-partner-specific version of NeCLAS (labeled *NoPair*), which comprises the same chemical features, but omits any nanoparticle information (Supplementary Information, section 4).

Performances for protein-nanoparticle predictions are shown in Fig. 2. Since the training of machine learning methods will be influenced by its training set and initial state, for NeCLAS and *NoPair* we show a distribution obtained from 250 different conditions, namely 25 different training and validation sets, each with 10 different initial sets of model weights.

**Fig. 2.**
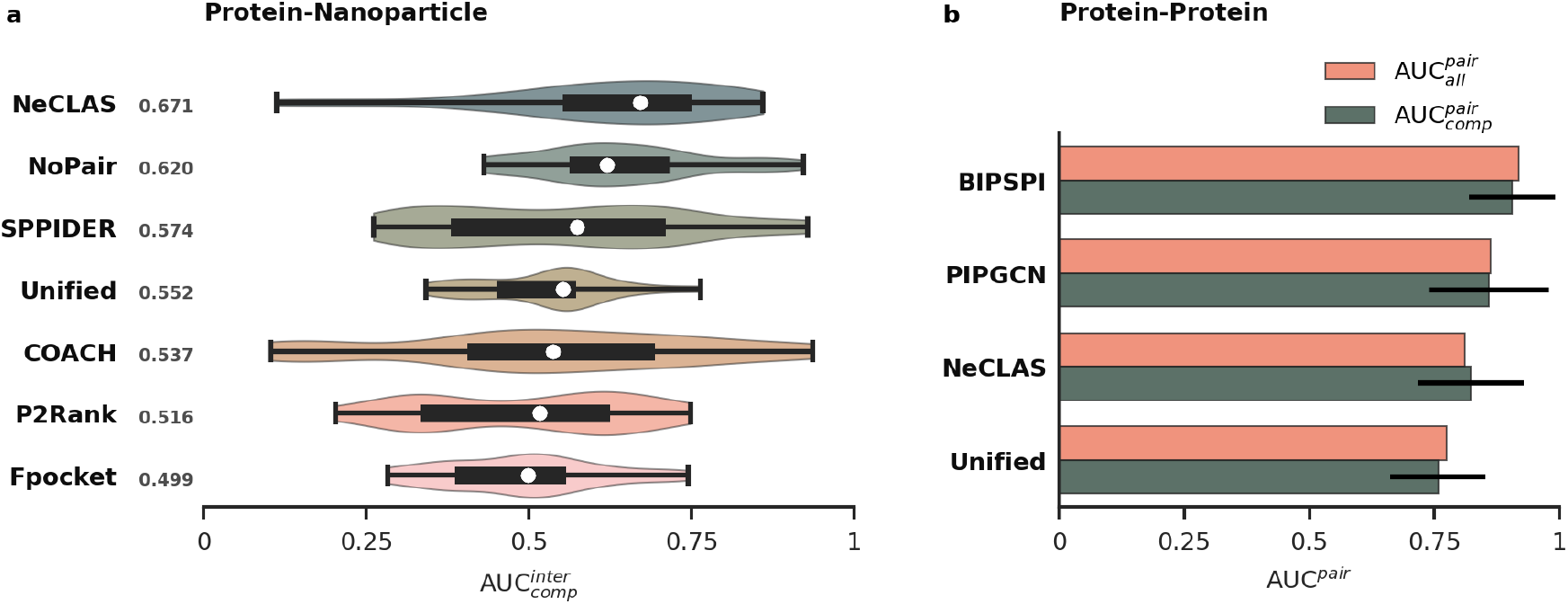
Predictive performances of different methods. **a**, Distribution of the prediction performance for protein-nanoparticle interface interactions. Median is marked as a white circle and reported as a number near the method name. The thick black bar shows the 1st-3rd interquartile range. NeCLAS and *NoPair* distributions are obtained by computing the median of each pair over 250 independent predictions. **b**, Performances of different methods for protein-protein pairwise interactions. Black lines indicate the standard deviation of complex-wise predictions.

NeCLAS outperforms all competing methods, with a median 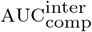 of 0.671, and the highest median 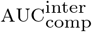 among all the 250 NeCLAS training conditions of 0.765. The closest existing method has a median 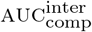 of 0.574. *NoPair* performs slightly worse (median 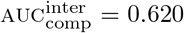), still surpassing the benchmark methods, which suggests that there is a performance benefit in including the representation of the partner molecules. It is important to note that the dataset is challenging for all the methods, and given the small number of nanoparticle-protein pairs it is not surprising to find that NeCLAS’s long tail is only due to two complexes. These results suggest that adding structural information to the validation set from similar nanoparticles may help generalize the stopping criteria of the neural network and improve NeCLAS performance.

As previously discussed, the structural homology between the training and testing datasets is a persistent issue in protein interaction predictions, leading to overly optimistic error estimates. Garcia *et al*. [20] considered this problem in the context of protein-protein interactions. They evaluated pairwise interactions by removing pairs of proteins that share both SCOP (Structural Classification of Proteins) families or a single SCOP family [47, 48]. Here, these criteria are already met, since the chosen nanoparticles are not structurally homologous to any proteins. However, we investigated the effect of an even stricter criterion by removing all proteins from the training and validation sets that share a single SCOP family with any of the proteins in our protein-nanoparticle test set. This test causes a negligible change in performance (median 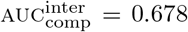, see Supplementary Information, section 4), showing no general effect due to the ablation.

Finally, we show that, despite its high degree of generality, NeCLAS achieves proteinprotein pairwise prediction performance that is competitive to state-of-the-art protein-specific methods (Fig. 2b). Our model is comparable (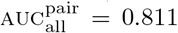 and median 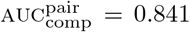) with PIPGCN [23] (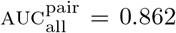 and median 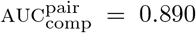 in its optimal configuration), and falls just below BIPSPI [20] (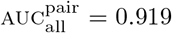 and median 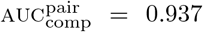), the currently leading methods in protein pairwise interaction predictions. Unlike competing methods, however, NeCLAS achieves these results while maintaining generality and omitting protein-specific (*e.g*., sequence features) information from the feature set. In addition, NeCLAS has minimal computational requirements, with a preprocessing time for DBD of approximately two hours on a consumer CPU (two orders of magnitude faster than Unified) and 1/10th the training time of PIPGCN on identical hardware.

To illustrate the specific potential of NeCLAS, we analyze in detail its performance on three system: nanoparticle tweezer and 14-3-3*σ* protein, carbon-based nanoparticle with amyloid fibrils, and organic quantum dots in water.

### Molecular Tweezers

Supramolecular ligands, such as molecular tweezers, represent a promising way to modulate protein functions. They can be artificially synthesized with unique properties and recognition profile towards amino acids and peptides, with ability to bind to specific sites. Specifically, the interactions between a 14-3-3*σ* protein and the lysine-specific molecular tweezers shown in Fig. 3 have been characterized in detail both experimentally and computationally [49]. Therefore, they serve as an ideal test case for pairwise interaction prediction models.

**Fig. 3.**
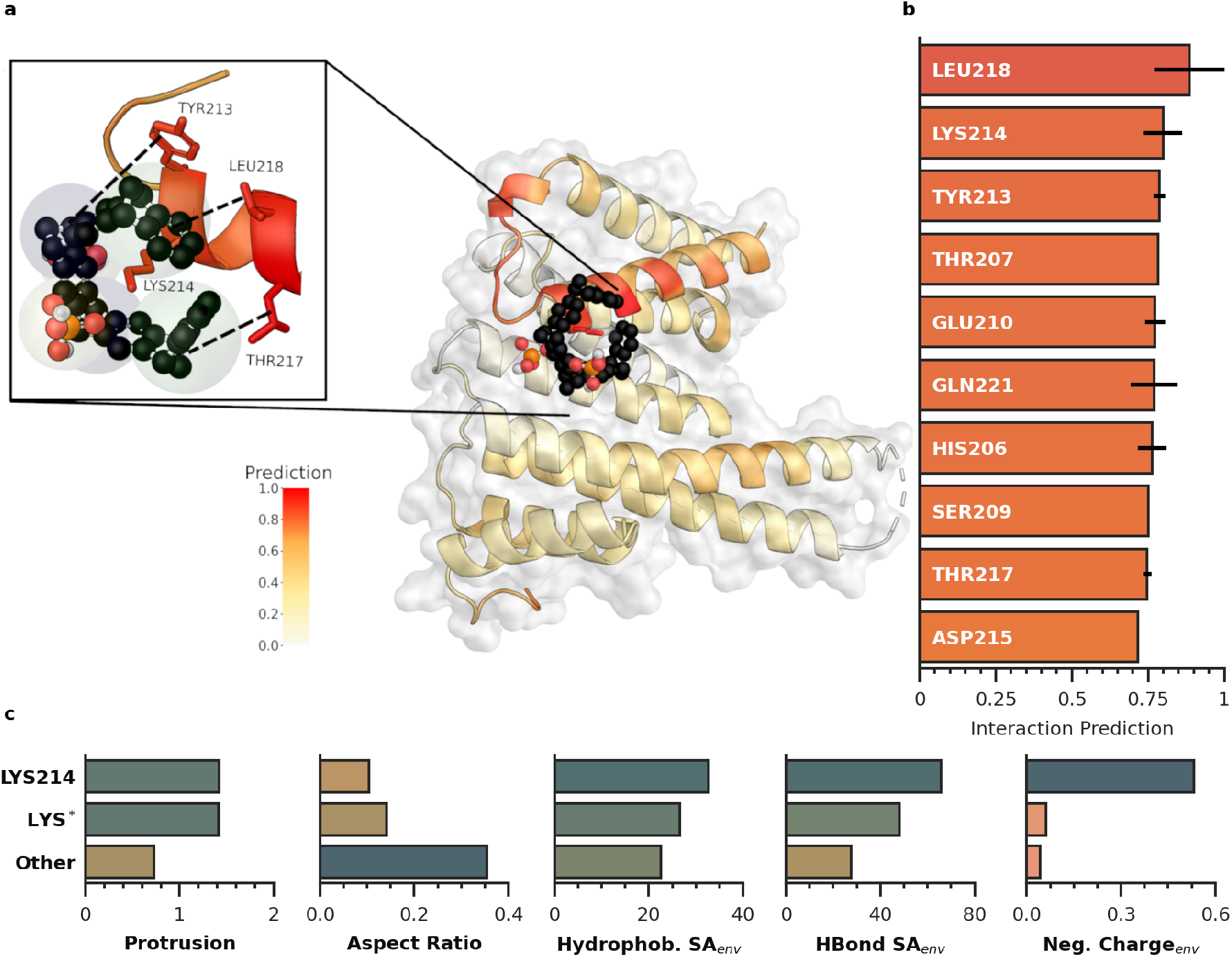
Interactions between molecular tweezers and the 14-3-3*σ* protein. **a**, Visual representation of the interaction location based on NeCLAS prediction; cutout highlights the lysine residue (Lys214) and the surrounding hydrophobic pocket. **b**, Top 10 interacting residues according to NeCLAS predictions. Black lines indicate standard deviation. All atoms in a single CG site share the same prediction. **c**, Comparison of selected features for binding lysine Lys214), probable but non-binding lysines (Lys*, as defined by Bier *et al*.), and all the other residues. Protrusion and aspect ratio are features of the individual amino acid sites (deterministic), while the last three histograms refer to environmentally weighted features.

NeCLAS predictions (Fig. 3a,b) match the findings reported by Bier *et al*., indicating the critical role played by Lys214, as well as Leu218, Tyr213, and Thr217, which form a hydrophobic binding pocket, and Glu210 and Gln221, which provide hydrogen bond stabilization. Bier *et al*. also derived a few general principles characterizing the active binding site (Lys214) leveraging the fact that the protein has four other energetically possible, but non-binding, lysine residues, (Lys* in the following) and that several properties differentiate Lys214 from the other residues.

First, they determined that Lys214 and Lys* residues are more energetically likely to bind due to their protruding carbon side chains, a characteristic that is matched by the higher value of the protrusion index [36] and elongated structure we observe in our model for Lys* residues compared to other residues (Fig. 3c). Additionally, Bier *et al*. suggested that the difference between Lys214 and the other Lys* residues is caused by the nearby hydrophobic binding pocket and small number of close positively charged functional groups, which destabilize the nanoparticle by forming external ion pairs between the nanoparticle phosphate groups and surrounding cations. These characteristics are matched in our model by several environmental features of Lys214 that capture the effect of neighboring atoms. The hydrophobic pocket for Lys214, depicted by the total surface area of surrounding hydrophobic groups [33] and total surface area of surrounding hydrogen bonding groups [34], shows higher values than all other residues. Additionally, the environmentally weighed charge, shows that Lys214 is surrounded by significantly more negatively charged atoms than other Lys* residues.

Beside displaying the predictive capabilities of NeCLAS, this comparison highlights the importance of spatially weighted physicochemical properties to fully describe nanoscale interactions. For proteins, we have shown that the same information can be encoded using only spatial features [50]. However, the much larger chemical and physical variety of nanoparticles properties requires a more nuanced approach, like the one used here, as nanoparticles with identical, or very similar, structures, but different properties are possible. As an additional benefit, these features provide a modicum of interpretability of the model’s predictions, which is harder to obtained when using more abstract properties.

### Bacterial Amyloid Fibrils

Interactions between nanoscale structures can exhibit extremely complex, high-dimensional free energy surfaces, which are the product of dynamic molecular constraints and entropic factors. Molecular dynamics (MD) simulations can be used to model these high dynamic processes evolving across relatively short time scales.

Characterization of these interactions via ML is challenging and the comparison between the results obtained with these two approaches is complex, especially since MD generates ensemble distributions of conformations. While datasets, like DBD version 5 used here, do not contain all the information that play a role in shaping the free energy landscape and the dynamics of a nanoscale system, ultimately, ML and MD are both (different) representations of the same physical system.

In order to further assess NeCLAS’s ability to predict nanoscale interactions, we chose to analyze the interactions between phenol-soluble modulin (PSM*α*1) peptides and graphene quantum dots (GQDs) [51]. We have previously shown that GQD nanoparticles act as biofilm dissolving agents due to their interactions with PSM*α*1, a key constituent of *Staphylococcus aureus* biofilm matrix, that assemble into amyloid fibers (Fig. 4a).

**Fig. 4.**
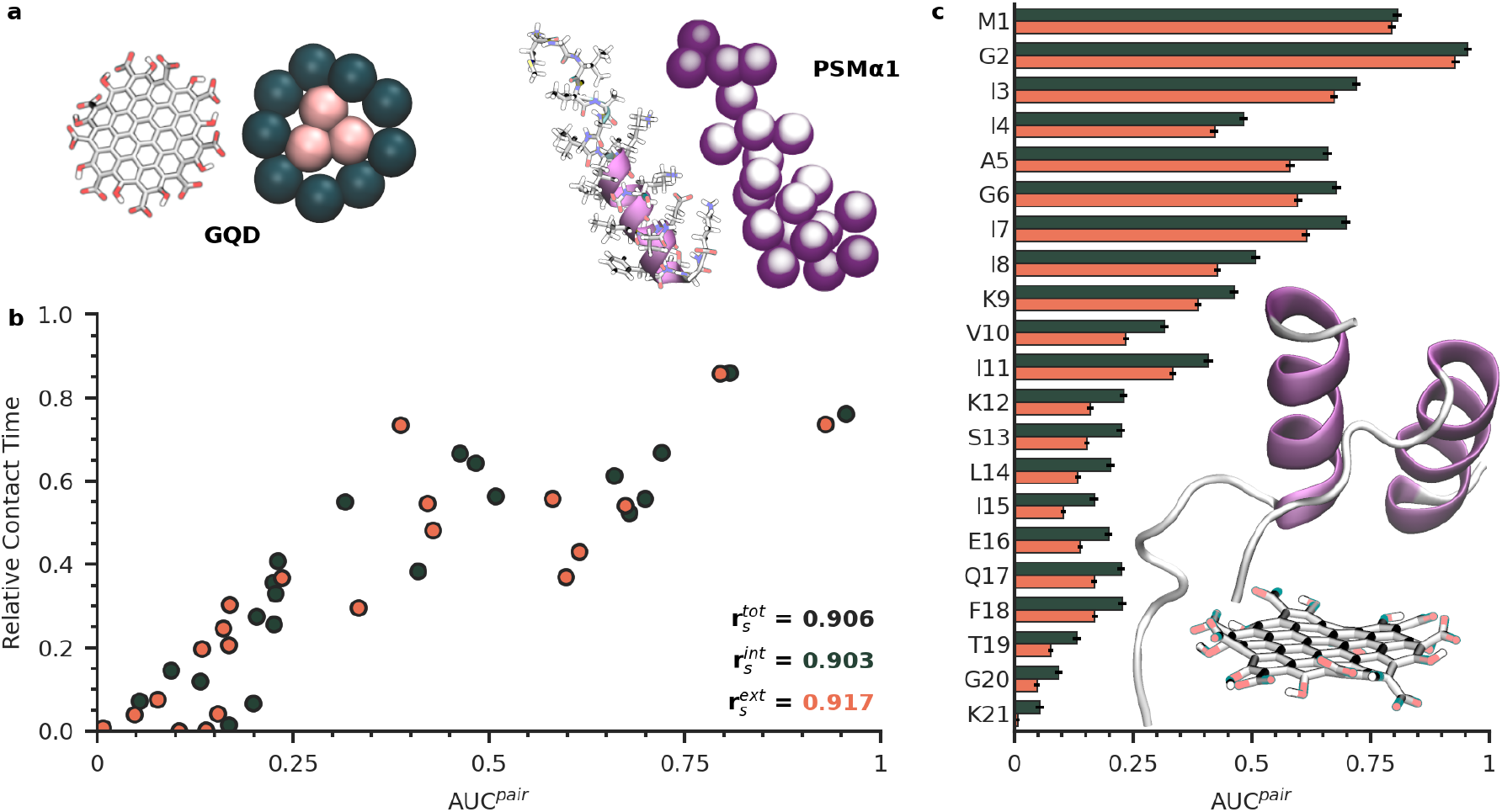
Interactions of PSM*α*1 1 and graphene quantum dot. **a**, All-atom and CG representation of the GQD (circumcoronene with alternating hydroxyl and carboxyl groups at the edges) and the PSM*α*1 1 peptide (5KHB). **b**, Relation between MD contact time and predicted pairwise interactions between the PSM*α*1 residues and the GQD sites. Spearman correlation 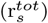 for all, interior only 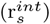 and exterior only 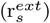 sites is reported. **c**, Average interaction prediction between residues from PSM*α*1 1 and GQD interior (green) and exterior (red) units, along with a snapshot of interactions observed during simulation. Standard deviation shown as black line.

To evaluate the ability of our model to predict the interactions between PSM*α*1 and GQDs, we compared the interaction probabilities obtained from NeCLAS with the contact times (*i.e*., the time two CG sites spent within a 1 nm-distance) during molecular dynamics (MD) simulations of the system composed of GQDs and PSM*α*1 as reported in Fig. 4b. The figure shows that the confidence of the predicted interactions is generally correlated with contact times (Spearman coefficient, 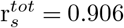). This trend is not simply due to strong interactions with the hydrophilic charged groups at the edges of the GQD, but rather due to a complex interplay of different chemical properties. To support this point, we chose to represent the GQD with 12 sites, which the CG procedure consistently separated in two distinct classes of nearly identical internal and external sites. The former contains only matrix carbon atoms while the latter includes the edge carbons and outer functional groups, thus allowing to separate the contributions of the two regions. The predictions for these subsets still show a high correlation 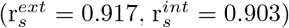 with the contact time, despite the different properties of these two types of sites. While internal hydrophobic sites generally have marginally lower predictions and shorter contact times, there is no separation between the two groups.

Finally, by analyzing the predictions for individual amino acids (Fig. 4c), we also confirm the importance of the N-terminal residues (which have the highest interaction probability), likely due to the GQD’s negative charge (dissociated carboxylic groups), in agreement with the previous observations by Wang *et al*. [51]. Of note, these conclusions are not dependent overall on the specific definition of contact time, and hold even when different definitions are used (Supplementary Information, section 5).

### Organic quantum dots

As a last example of NeCLAS potential, we discuss the ability to use pairwise interaction predictions to inform atomistic models, for example, to generate realistic conformation distributions or to evaluate the aggregation of multiple nanoparticles. In doing so, we further test the potential applications of our code. In these cases, a variety of factors (*e.g*., thermal energy, solvent effects, entropic contributions), need to be taken into account, which requires additional assumptions and data.

Furthermore, the scalar predictions of binary classification models cannot be directly interpreted as a measure of interaction strength, and they are more readily conceptualized as model confidence. However, here, we consider the probability of interaction as being proportional to the strength of interaction, as we expect a well-informed model to classify weakly interacting pairs as less likely to interact. Under this assumption, we use NeCLAS predictions to tune the intensity of intermolecular forces of different GQDs in water to study their propensity to form aggregates. Previously, using *all-atom* molecular dynamics, we have reported on the effect of the composition of edge groups present on GQDs and their tendency to aggregate in water [52].

Here, we study three types of GQDs: one terminated with hydroxyl (g3oh), another with formyl (g3cho) groups, and the last one with an alternating 2:1 ratio of hydroxyl and cysteine groups (6C-g3oh). These nanoparticles, with sizes between 1.5 and 2 nm, were chosen as hydrophobic and hydrophilic forces are generally comparable, whereas for bigger structures, hydrophobic forces and water entropic exclusion increasingly dominate their interactions. Pairwise NeCLAS predictions (AUC^pair^) were converted to an intermolecular potential by using a tunable repulsive-core potential that uses identical parameters for all the sites of the GQDs, except for the single value (NeCLAS prediction) that varies between 0 and 1. This step allows converting NeCLAS predictions into a physically-meaningful potential while reducing instabilities for weak interactions (Fig. 5a,d).

**Fig. 5.**
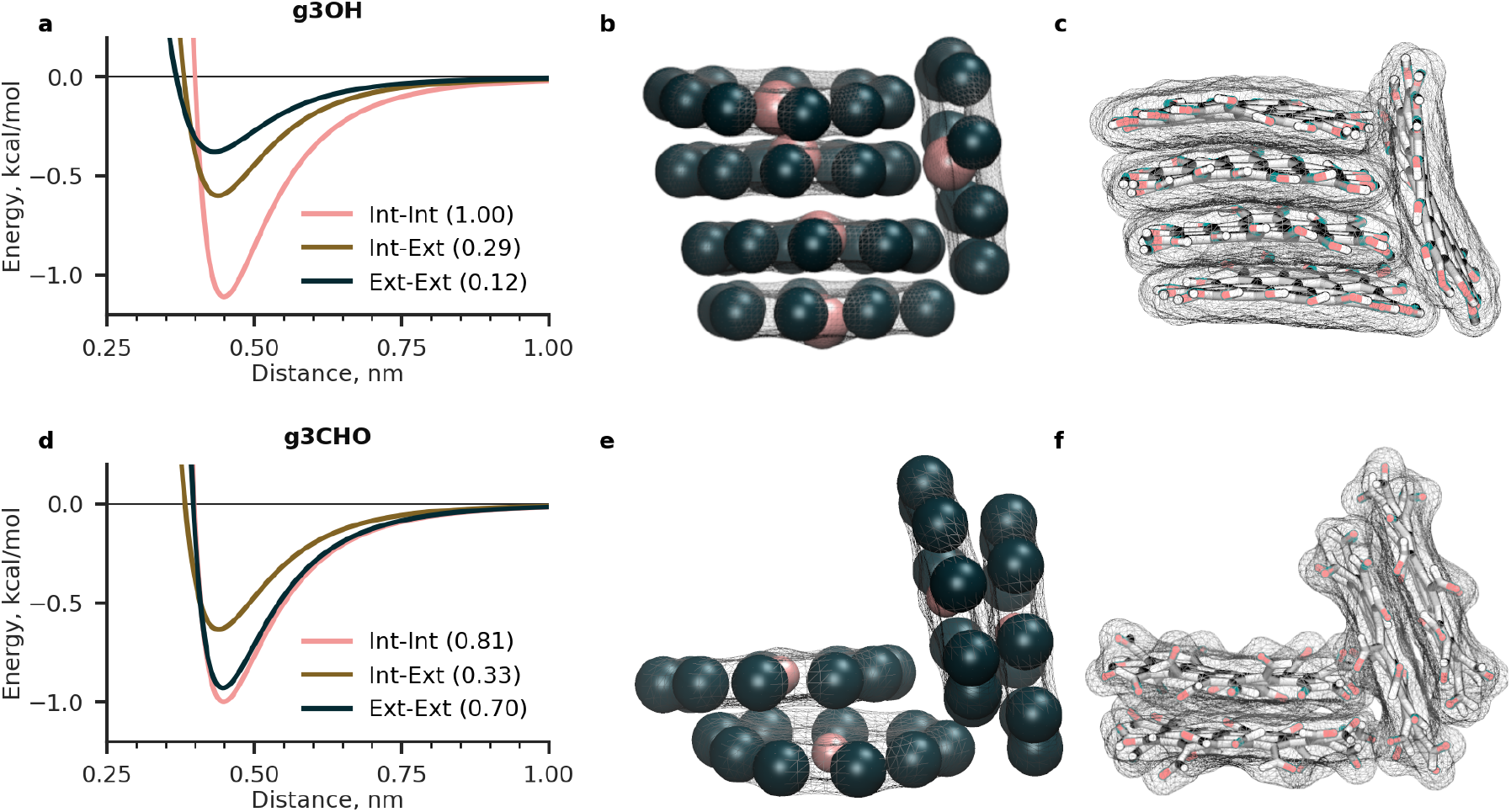
Predicted and simulated interactions of graphene quantum dots. **a**, pairwise interaction potential from NeCLAS mean predictions (in parentheses) for g3oh CG sites. **b**, Snapshot of g3oh from CG simulations modeled using NeCLAS predictions. **c**, Snapshot of all-atom g3oh simulations. **d**, pairwise interaction potential from NeCLAS mean predictions (in parentheses) for g3cho CG sites. **e**, Snapshot of g3cho from CG simulations modeled using NeCLAS predictions. **f**, Snapshot of all-atom g3cho simulations.

Using these potentials, we simulated the dynamics of a small number of GQDs randomly placed in a periodic system, using coarse-grained MD. For g3oh and g3cho we observed the rapid formation of aggregates with structures that closely resemble the ones observed in allatom molecular dynamics [52] (Fig. 5). Indeed, we detect both close and parallel stacking of the structures (see [52] for definition), and a similar lateral shift between consecutive stacking planes. 6C-g3oh, however, did not aggregate, again in agreement with AA simulations and the experimental high solubility at pH 7 [53]. The overall match between CG and all-atom simulations, albeit qualitative, was obtained without fine-tuning of the other parameters of the CG potential, as such optimizations would obscure the contributions of NeCLAS. Better agreements can be expected if additional optimizations are performed.

## Discussion

The previous results show that NeCLAS can predict protein-protein and protein-nanoparticle pairwise interactions, and can be extended to predict nanoparticle-nanoparticle systems, while using properties that are general to all molecules. The quality of NeCLAS predictions hinges on two main aspects: the simplification operated through the CG representation and the use of environmental features that capture the chemical neighborhood information.

The CG representation reduces physical (*e.g*., thermal vibration), observational (*e.g*., experimental and numerical error), and statistical (*e.g*., sampling size) uncertainties in the data, allowing an efficient and robust model training. This approach is not needed for all types of problems, but it is critical for data-limited applications, which often occur in chemistry. Furthermore, coarse-graining lowers the computational requirements, with the potential for applications to larger systems, while allowing to reconstruct atomistic information if needed [54].

However, a CG representation causes the loss of local atomistic properties, which are not adequately captured through averaging. Here, we express the local spatial distribution of each property using various statistics (*e.g*., standard deviation), which provides a more nuanced characterization of the target distribution. Yet, this approach does not solve the problem of long-range nanoscale interactions, which tend to decay quickly in solvents with high relative permittivity, but can still play a critical role in long-range organization (*e.g*., tertiary structures in proteins). To capture these interactions, we computed properties by weighting atomic characteristic of neighboring atoms (part of different CG sites) by their distance from a given site, which simultaneously embeds information about both properties and positions of the individual atoms.

The local atomic arrangement is critical to capture interactions at this scale, and we have shown that, for proteins, it is an equivalent surrogate for describing physicochemical properties [50]. Differently from previous works, we did not find that structural information alone can provide a general enough description for both particles and nanoparticles. Even if structural information is sufficient for some classes of molecules (*e.g*., proteins), in which the limited chemical variety results in a correlation between atomic species and spatial organization, they become insufficient to distinguish wider classes of systems. One such examples, are fullerenes, which can gain different amounts of charge while in water without a relevant change in structure [55]. NeCLAS predicts a marked difference (71% increase) in the interactions between a fullerene organizing protein (PDB: 5ET3) [56] and a neutral C60 fullerene (as it is commonly modelled) or a negatively charged one (average experimental charge of −2 elementary charges).

NeCLAS also addresses the effect of data symmetries on model performance and reliability. Many interaction-prediction methods use ensemble-based models (*e.g*., XGBoost) [20, 22] or dense neural networks [44] to predict interaction interfaces or pairwise interactions. However, both of these methods are permutation variant, adding artificial ordering to a problem that is inherently unordered. In doing so, these methods violate the guiding principle that subjective ordering has no bearing on the behavior of a physical system. Further, these models produce unstable prediction results, as the hypothesis space of an over-parameterized model may contain many permutation variant functions that fit the training data (Supporting Section 4). NeCLAS avoids this problem by using a permutation invariant neural network inspired by the *DeepSets* [57] architecture. Thanks to a combination of weight sharing and maxpooling, our model is agnostic to residue ordering, and empirically outperforms comparable permutation variant methods while utilizing significantly fewer weights.

Despite the positive results of NeCLAS, there are still some aspects that deserve further work. To operate on an unconstrained chemical space of nanoparticles, NeCLAS will require a more diverse sample of atom types and solvents. For example, NeCLAS (and other methods) tacitly assume that most species of interest are largely soluble in water. Under these conditions, many of the forces that govern protein complexes are also present in interactions between the proteins and nanoparticles [43], as water solubility strongly limits the chemical properties of exposed nanoparticle surfaces. Different solvent (*e.g*., polymeric hostguest systems) not only have different ability to stabilize ionic groups or form hydrogen bonds, but also a different propensity than water to solvate species based on their size. A more varied dataset can also potentially highlight the need for a more accurate or diverse representation of chemical variety or large scale structures.

All the above limitations, however, can only be addressed by the availability of curated data that go beyond protein-protein interactions. As more structural information and databases for nanoscale species emerge, we expect that this approach will prove to be a valuable technique for operating across new molecule types and biological interaction problems.

## Methods

### Coarse-Grained representation

Coarsegrained representations of nanoparticles and proteins were created using the neural gas algorithm [31], which uses a Hebbian network to match the mass target distribution. Details about convergence, hyperparameters choice and robustness are discussed in the Supporting Information, section 1.

### Physicochemical Features

We used 80 features to describe the local chemistry (local descriptors) of each CG site and 400 features to describe the chemical neighborhood (environmental descriptors). The local chemistry was captured by using mass, charge, relative accessible surface area [38], depth [35], protrusion [36], CPSA descriptors [33], CPSA hydrogen bonding descriptors [34], pocket propensity, and mass-weighted WHIM [37]. Depth and protrusion values were computed for each atom in the residue and for depth protrusion, charge, and mass a feature set was generated by taking the minimum, maximum, mean, sum, and standard deviation of the values. Charges were computed by processing proteins with the PDB2PQR package [58] using the AMBER force field [59], and computed for nanoparticles with the Gasteiger method [60].

Environmental descriptors were computed using five different sets of environmental [41] functions for each feature to compute 400 properties. See section 2 in the Supporting Information for additional details.

### Protein-Protein Dataset

For protein-protein pairwise predictions, we used the DBD (version 5) [24]. Unbound proteins were used for feature generation, and bound proteins were used to compute the ground-truth pairwise interactions. For each complex, we consider all combinations of residues between the two proteins. Due to the severe class imbalance (positive sample rate of 0.136%), we downsampled our training data so that there is one positive example for every three negative examples, providing approximately 83,000 pairwise interactions (different validation splits cause the exact value to change). We did not alter the imbalance from the testing set to avoid biasing our data.

### Nanoparticle-Protein Dataset

For the protein-nanoparticle interactions, we used the recent collection of crystallographic data by Costanzo’ *et al*. [43]. This collection contains approximately 40 unique structures; however, we removed files containing duplicate interactions or incomplete information, leaving 21 unique protein-nanoparticle pairs (see section 3 in the Supporting Information). Unbound proteins structures were taken from the RCSB database [61], while unbound nanoparticle structures were generated by relaxing the bounded configuration with the MMFF94 force field [62] in the absence of the protein.

### ML procedure

We implemented a custom, alternating structure of permutation variant networks and maxpool to derive a generalized, non-linear, permutation-invariant network. For protein-protein pairwise prediction, we performed a leave-one-out cross validation. For proteinnanoparticle pairwise predictions, we trained on the entire protein-protein dataset (downsampled as described above), and used the proteinnanoparticle dataset for testing. Interface interaction predictions were obtained from pairwise predictions using a scoring function [20]. All predictions were smoothed [20, 22].

For each pair of CG sites, the local and environmental residue features of both sites were concatenated to create a single input feature to the model. Two sites *A, B* are considered to be interacting if and only if a heavy atom from site *A* falls within 0.6 nm of a heavy atom of site *B*. When conversion between CG predictions and protein residues was needed, we assigned each heavy atom a prediction score equal to that of its corresponding CG site and computed the residue prediction as the mean of all its constituent atoms (excluding hydrogen). Additional details can be found in the Supplementary information, section 4.

### Molecular Dynamics

In the CG simulations, 8 particles were randomly placed in a box, and after an energy minimization, the system was run for 5 ns in a canonical ensemble at 300 K. Snapshots were taken from the last 500 ps of each run. All-atom simulation parameters, protocol, and force filed are described in detail elsewhere [52]. Additional details can be found in Supporting Information, section 5.

### Data processing and visualization

Data were processed with custom Python code using the NumPy library [63]. Molecular images were generated using the Visual molecular dynamics code [64] and Pymol [65]. Plots were created using Matplotlib [66].

## Additional Information

### Data availability

Additional data will be available at DeepBlue (https://deepblue.lib.umich.edu/TBD), an open and permanent data repository maintained by the University of Michigan.

### Code availability

The code used in this work together with the relative documentation will be available on Code Ocean (https://codeocean.com/TBD).

## Acknowledgments

M.R. thanks Clayton Scott for insightful feedback and discussions. P.E. thanks Chloe Luyet for the help with the all-atom simulation of 6C-g3oh. The work was supported by the BlueSky Initiative, The University of Michigan College of Engineering, PI: A. Violi. We acknowledge Advanced Research Computing, a division of Information and Technology Services at the University of Michigan, for computational resources and services provided for the research.

## Author contributions

A.V. and P.E. conceived and supervised the project. J.S. and P.E. conceived chemical features and representations.

M.R. and J.S. designed, trained, and tested the machine learning models. J.S. designed experiments and created the database. P.E. designed and ran the MD simulations. All authors read, revised and approved the manuscript.

## References

[1] Ghosh, G. & Panicker, L. Protein–nanoparticle interactions and a new insight. Soft Matter 17 (14), 3855–3875 (2021).

[2] Russ, K. A. et al. C _60_fullerene localization and membrane interactions in RAW 264.7 immortalized mouse macrophages. Nanoscale 8 (7), 4134–4144 (2016).

[3] Liu, C. et al. Predicting the Time of Entry of Nanoparticles in Lipid Membranes. ACS Nano 13 (9), 10221–10232 (2019).

[4] Auría-Soro, C. et al. Interactions of Nanoparticles and Biosystems: Microenvironment of Nanoparticles and Biomolecules in Nanomedicine. Nanomaterials 9 (10), 1365 (2019).

[5] Cha, S.-H. et al. Shape-Dependent Biomimetic Inhibition of Enzyme by Nanoparticles and Their Antibacterial Activity. ACS Nano 9 (9), 9097–9105 (2015).

[6] Salvati, A. et al. Transferrin-functionalized nanoparticles lose their targeting capabilities when a biomolecule corona adsorbs on the surface. Nature Nanotechnology 8 (2”), 137–143 (2013).

[7] Abarca-Cabrera, L., Fraga-García, P. & Berensmeier, S. Bio-nano interactions: Binding proteins, polysaccharides, lipids and nucleic acids onto magnetic nanoparticles. Biomaterials Research 25 (1), 12 (2021).

[8] Adcock, S. A. & McCammon, J. A. Molecular dynamics: Survey of methods for simulating the activity of proteins. Chemical Reviews 106 (5), 1589–1615 (2006).

[9] Aubin-Tam, M.-E. & Hamad-Schifferli, K. Structure and function of nanoparticle–protein conjugates. Biomedical Materials 3 (3), 034001 (2008).

[10] Ge, C. et al. Binding of blood proteins to carbon nanotubes reduces cytotoxicity. Proceedings of the National Academy of Sciences 108 (41), 16968–16973 (2011).

[11] Perilla, J. R. et al. Molecular dynamics simulations of large macromolecular complexes. Current Opinion in Structural Biology 31, 64–74 (2015).

[12] Vilanova, O. et al. Understanding the kinetics of protein–nanoparticle corona formation. ACS Nano 10 (12), 10842–10850 (2016).

[13] Schneidman-Duhovny, D., Inbar, Y., Nussinov, R. & Wolfson, H. J. PatchDock and SymmDock: Servers for rigid and symmetric docking. Nucleic Acids Research 33 (Web Server), W363–W367 (2005).

[14] Pierce, B. G. et al. ZDOCK server: Interactive docking prediction of protein-protein complexes and symmetric multimers. Bioinformatics 30 (12), 1771–1773 (2014).

[15] Yan, Y., Tao, H., He, J. & Huang, S.-Y. The HDOCK server for integrated protein–protein docking. Nature Protocols 15 (5), 1829–1852 (2020).

[16] Lim, S. et al. A review on compound-protein interaction prediction methods: Data, format, representation and model. Computational and Structural Biotechnology Journal 19, 1541–1556 (2021).

[17] Krivaák, R. & Hoksza, D. P2Rank: Machine learning based tool for rapid and accurate prediction of ligand binding sites from protein structure. Journal of Cheminformatics (1), 39 (2018).

[18] Gainza, P. et al. Deciphering interaction fingerprints from protein molecular surfaces using geometric deep learning. Nature Methods 17 (2), 184–192 (2020).

[19] Townshend, R., Bedi, R., Suriana, P. & Dror, R. End-to-End Learning on 3D Protein Structure for Interface Prediction. Neural Information Processing Systems (33), 15642–15651 (2019).

[20] Sanchez-Garcia, R., Sorzano, C., Carazo, J. M. & Segura, J. Bipspi: a method for the prediction of partner-specific protein-protein interfaces. Bioinformatics 35 (3), 470–477 (2019).

[21] Dai, B. & Bailey-Kellogg, C. Protein interaction interface region prediction by geometric deep learning. Bioinformatics 37 (17), 2580–2588 (2021).

[22] ul Amir Afsar Minhas, F., Geiss, B. J. & Ben-Hur, A. Pairpred: Partner-specific prediction of interacting residues from sequence and structure. Proteins 82, 1142–1155 (2014).

[23] Fout, A., Byrd, J., Shariat, B. & Ben-Hur, A. Protein interface prediction using graph convolutional networks. NIPS 30 (2017).

[24] Vreven, T. et al. Updates to the integrated protein–protein interaction benchmarks: Docking benchmark version 5 and affinity benchmark version 2. Journal of Molecular Biology 427 (19), 3031–3041 (2015).

[25] Monopoli, M. P., Aberg, C., Salvati, A. & Dawson, K. A. Biomolecular coronas provide the biological identity of nanosized materials. Nature Nanotechnology 7 (12), 779–786 (2012).

[26] Saptarshi, S. R., Duschl, A. & Lopata, A. L. Interaction of nanoparticles with proteins: Relation to bio-reactivity of the nanoparticle. Journal of Nanobiotechnology 11 (1), 26 (2013).

[27] Zuo, G., Kang, S.-g., Xiu, P., Zhao, Y. & Zhou, R. Interactions between proteins and carbon-based nanoparticles: Exploring the origin of nanotoxicity at the molecular level. Small 9 (9-10), 1546–1556 (2013).

[28] Findlay, M. R., Freitas, D. N., Mobed-Miremadi, M. & Wheeler, K. E. Machine learning provides predictive analysis into silver nanoparticle protein corona formation from physicochemical properties. Environmental Science: Nano 5 (1), 64–71 (2018).

[29] Ban, Z. et al. Machine learning predicts the functional composition of the protein corona and the cellular recognition of nanoparticles. Proceedings of the National Academy of Sciences 117 (19), 10492–10499 (2020).

[30] Ouassil, N., Pinals, R. L., Bonis-O’Donnell, J. T. D., Wang, J. W. & Landry, M. P. Supervised learning model predicts protein adsorption to carbon nanotubes. Science Advances 8 (1), eabm0898 (2022).

[31] Martinetz, T., Berkovich, S. & Schulten, K. ‘Neural-gas’ network for vector quantization and its application to time-series prediction. IEEE Transactions on Neural Networks 4 (4), 558–569 (1993).

[32] Alex, J. M. et al. Calixarene-mediated assembly of a small antifungal protein. IUCrJ 6 (2), 238–247 (2019).

[33] Stanton, D. T. & Jurs, P. C. Development and use of charged partial surface area structural descriptors in computer-assisted quantitative structure-property relationship studies. Analytical Chemistry 62 (21), 2323–2329 (1990).

[34] Stanton, D. T., Egolf, L. M., Jurs, P. C. & Hicks, M. G. Computer-assisted prediction of normal boiling points of pyrans and pyrroles. Journal of Chemical Information and Computer Sciences 32 (4), 306–316 (1992).

[35] Sanner, M. F., Olson, A. J. & Spehner, J.-C. Reduced surface: An efficient way to compute molecular surfaces. Biopolymers 38 (3), 305–320 (1996).

[36] Pintar, A., Carugo, O. & Pongor, S. CX, an algorithm that identifies protruding atoms in proteins. Bioinformatics 18 (7), 980–984 (2002).

[37] Todeschini, R. & Gramatica, P. The Whim Theory: New 3D Molecular Descriptors for Qsar in Environmental Modelling. SAR and QSAR in Environmental Research 7 (1-4), 89–115 (1997).

[38] Kawabata, T. Detection of multiscale pockets on protein surfaces using mathematical morphology. Proteins: Structure, Function, and Bioinformatics 78 (5), 1195–1211 (2010).

[39] Porollo, A. & Meller, J. Prediction-based fingerprints of protein-protein interactions. Proteins: Structure, Function, and Bioinformatics 66 (3), 630–645 (2006).

[40] Behler, J. Atom-centered symmetry functions for constructing high-dimensional neural network potentials. The Journal of Chemical Physics 134 (7), 074106 (2011).

[41] Gastegger, M., Schwiedrzik, L., Bittermann, M., Berzsenyi, F. & Marquetanda), P. wacsf—weighted atom-centered symmetry functions as descriptors in machine learning potentials. J. Chem. Phys. 148 (2018).

[42] Clark, J. J., Orban, Z. J. & Carlson, H. A. Predicting binding sites from unbound versus bound protein structures. Scientific Reports 10 (1), 243–252 (2020).

[43] Costanzo, L. D. & Geremia, S. Atomic details of carbon-based nanomolecules interacting with proteins. Molecules 25 (15), 3555 (2020).

[44] Cha, M. et al. Unifying structural descriptors for biological and bioinspired nanoscale complexes. Nature Computational Science 2, 243–252 (2022).

[45] Yang, J., Roy, A. & Zhang, Y. Protein-ligand binding site recognition using complementary binding-specific substructure comparison and sequence profile alignment. Bioinformatics (Oxford, England) 29 (20), 2588–2595 (2013).

[46] Le Guilloux, V., Schmidtke, P. & Tuffery, P. Fpocket: An open source platform for ligand pocket detection. BMC Bioinformatics 10 (1), 168 (2009).

[47] Andreeva, A., Howorth, D., Chothia, C., Kulesha, E. & Murzin, A. G. SCOP2 prototype: a new approach to protein structure mining. Nucleic Acids Research 42 (D1), D310–D314 (2013).

[48] Andreeva, A., Kulesha, E., Gough, J. & Murzin, A. G. The SCOP database in 2020: expanded classification of representative family and superfamily domains of known protein structures. Nucleic Acids Research 48 (D1), D376–D382 (2019).

[49] Bier, D. et al. Molecular tweezers modulate 14-3-3 protein–protein interactions. Nature Chemistry 234–239 (2013).

[50] Baranwal, M. et al. Struct2graph: A graph attention network for structure based predictions of protein-protein interactions. bioRxiv (2020).

[51] Wang, Y. et al. Anti-Biofilm Activity of Graphene Quantum Dots via Self-Assembly with Bacterial Amyloid Proteins. ACS Nano 13 (4), 4278–4289 (2019).

[52] Elvati, P., Baumeister, E. & Violi, A. Graphene quantum dots: effect of size, composition and curvature on their assembly. RSC Advances 29 (2017).

[53] Suzuki, N. et al. Chiral Graphene Quantum Dots. ACS Nano 10 (2), 1744–1755 (2016).

[54] Izvekov, S. & Voth, G. A. A multiscale coarse-graining method for biomolecular systems. The Journal of Physical Chemistry B 109 (7), 2469–2473 (2005).

[55] Deguchi, S., Alargova, R. G. & Tsujii, K. Stable Dispersions of Fullerenes, C60 and C70, in Water. Preparation and Characterization. Langmuir 17 (19), 6013–6017 (2001).

[56] Kim, K.-H. et al. Protein-directed self-assembly of a fullerene crystal. Nature Communications 7 (1), 11429 (2016).

[57] Zaheer, M. et al. Guyon, I. et al. (eds) Deep sets. (eds Guyon, I. et al.) Advances in Neural Information Processing Systems, Vol. 30 (Curran Associates, Inc., 2017).

[58] Dolinsky, T. J. et al. PDB2PQR: Expanding and upgrading automated preparation of biomolecular structures for molecular simulations. Nucleic Acids Research 35 (Web Server), W522–W525 (2007).

[59] Hornak, V. et al. Comparison of multiple Amber force fields and development of improved protein backbone parameters. Proteins: Structure, Function, and Bioinformatics 65 (3), 712–725 (2006).

[60] Gasteiger, J. & Marsili, M. A new model for calculating atomic charges in molecules. Tetrahedron Letters 19 (34), 3181–3184 (1978).

[61] Berman, H., Henrick, K. & Nakamura, H. Announcing the worldwide Protein Data Bank. Nature Structural & Molecular Biology 10 (12), 980–980 (2003).

[62] Halgren, T. A. Merck molecular force field. I. Basis, form, scope, parameterization, and performance of MMFF94. Journal of Computational Chemistry 17 (5-6), 490–519 (1996).

[63] Harris, C. R. et al. Array programming with NumPy. Nature 585 (7825), 357–362 (2020).

[64] Humphrey, W., Dalke, A. & Schulten, K. VMD: Visual molecular dynamics. Journal of Molecular Graphics 14 (1), 33–38 (1996).

[65] Schroödinger, L. & DeLano, W. Pymol.

[66] Hunter, J. D. Matplotlib: A 2d graphics environment. Computing in Science & Engineering 9 (3), 90–95 (2007).

