## Supplementary material for "From proteins to nanoparticles: domain-agnostic predictions of nanoscale interactions": Supplmenetary Information

### Supplementary Information

Jacob Saldinger[1], Matt Raymond[2], Paolo Elvati[3] and Angela Violi[1,2,3*]

[1]Chemical Engineering, University of Michigan, Street, Ann Arbor, 48109-2125, Michigan, USA.
[2]Electrical Engineering and Computer Science, University of Michigan, Street, Ann Arbor, 48109-2125, Michigan, USA.
[3]Mechanical Engineering, University of Michigan, Street, Ann Arbor, 48109-2125, Michigan, USA.

Contributing authors:;
;

### 1 Dimensionality Reduction

#### 1.1 Background

Our coarse-graining method is based on the neural gas algorithm from Martinez *et al.* [1]. Initial positions of N sites centers are assigned across the molecule with the *kmeans++* initialization algorithm [2]. The site's center positions are then iteratively updated to reach a final configuration.

For each iteration, an atom is stochastically selected with a probability according to a target property (mass). Each site is then numerically ranked, $k$, according to its proximity to the selected atom. Positions of each site are updated through,

$$R_i^{new} = R_i^{old} + \epsilon \exp\left[-k/\lambda\right](v - R_i^{old}) \tag{1}$$

where $R_i$ is the site position and $v$ is the coordinate of the selected atom. Parameters $\epsilon$ and $\lambda$ are adjusted according to,

$$p = p_o(p_s/p_o)^{\frac{s}{S}} \tag{2}$$

where $p$ is the parameter (either $\epsilon$ or $\lambda$), $p_o$ and $p_s$ are hyperparameters, s is the current step, and S is the total number of steps. After completion of all iterations, atoms are assigned to the closest site (euclidean distance). The number of iterations, hyperparameters, and the target property of mass are unchanged from previous implementations [3].

#### 1.2 Convergence

In this section, we briefly discuss the parameters used in our coarse-graining approach, their sensitivity, and their validation. The final values are reported below,

**Table S1** Hyperparameters used in the coarse-graining procedure.

| Variable | Value |
|---|---|
| $\epsilon_o$ | 0.3 |
| $\epsilon_s$ | 0.05 |
| $\lambda_o$ | 0.2 N |
| $\lambda_s$ | 0.01 |
| N | number of sites |
| S | 200 N |

To assess if these parameters are sufficient to obtain convergence on our dataset, we monitored the RMSD between the CG site centers and the final positions at each step of the coarse-graining process (Figure S1).

For all systems, the largest changes towards the final position occurs in the first half of the procedure, showing that additional steps or faster convergence

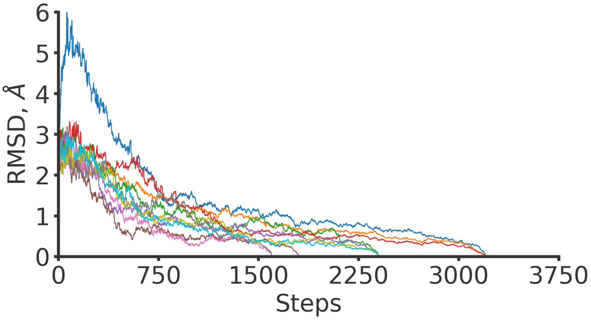

**Fig. S1 Convergence of coarse-graining procedure**, monitored as the RMSD of CG centers from final position *vs* number of steps in coarse-grained simulation. For clarity, only 10 nanoparticles are shown.

rates are not needed to reach the final positions. Note that the small decrease observed at the end of every trajectory is due to the fact that, by definition, the RMSD needs to equal zero at the final step, and RMSD is a non-negative metric.

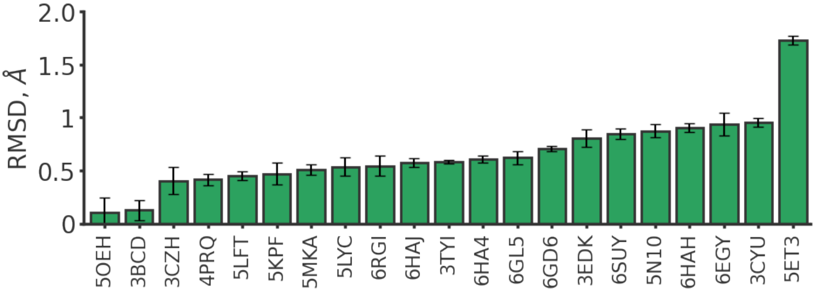

**Fig. S2 Stability of coarse-graining procedure**, defined as the mean RMSD of the final position after 10 distinct simulations for each nanoparticle in the dataset. More information about each system and x-axis labels can be found in Table S3.

#### 1.3 Stability

We also assess the stability of our predictions (Figure S2) to verify that multiple runs of the same coarse-graining do not give markedly different results. While the neural gas approach has been shown to produce more consistent solutions than other clustering algorithms [4], its stochasticity still can be a source of uncertainty in model predictions. Therefore, for all nanoparticles evaluated in our dataset, we compared the distances between the center of mass of CG sites across 10 replicate runs. In all but one cases, the average RMSD is less than 0.1 nm, which is less than the bond length of a carbon and hydrogen. This suggests that while some small stochastic fluctuations are possible, the sites are largely independent of starting position or random seed. We note that the only outlier is C60 (5ET3), which due to its symmetry can be represented with identical sites in different positions.

The only other parameter, used in the dimensionality reduction procedure, is the number of sites. This parameter does not affect the consistency of the convergence, but rather the stability of the neural network results. For protein coarse-graining, we set this value such that there are 7.5 atoms (excluding H atom) on average in each CG site, which corresponds approximately to the average amino acid size in our dataset. For nanoparticles, we select a similar size, however, slightly adjust the value to correspond to natural nanoparticle symmetries (see Figure 1 in main text). Exact numbers for each nanoparticle are given in section 3.

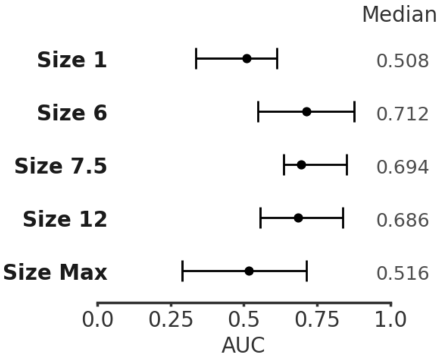

**Fig. S3 Distribution of $\text{AUC}^{\text{inter}}_{\text{comp}}$ using the nanoparticle dataset** for different number of atoms in CG site. Each size (y-axis) indicates the average number of atoms (excluding H atoms) in the CG sites; Max is used to indicate that the entire molecule is reduced to a single site. Circles show median value and lines show quartile ranges.

The quality of NeCLAS's protein-nanoparticle predictions as function of the nanoparticles CG site size is shown in Figure S3). Our findings suggest that our results are not highly sensitive to the number of sites, however, steep drop-offs in performance can occur if the CG site size is significantly larger or smaller than amino acids due to the scales of the interactions.

Finally, it is worth discussion the choice of mass as a target property. We chose this based on previous works [3] and expect it to be a good representation of properties for all atoms in our organic datasets since carbon, nitrogen, and oxygen have similar masses, sulfur is relatively rare in our molecules, and hydrogen contributes little to the final chemical descriptors. Still, it is worth noting that with other prediction problems and molecule types (*e.g.*, metal or ceramic nanoparticles) a different property such as charge, surface area, or electronegativity may better weight the atoms that contribute to an interaction.

### 2 Features

#### 2.1 Local properties

**Table S2** Descriptors used in model.

| Descriptor | Number | Details |
|---|---|---|
| CPSA | 29 | [5] |
| CPSA Hydrogen Bonding | 16 | [6] |
| WHIM | 14 | Mass-weighted [7] |
| Depth | 5 | Sum, min, max, std deviation, mean [8] |
| Protrusion | 5 | Sum, min, max, std deviation, mean [9] |
| Charge | 5 | Sum, min, max, std deviation, mean |
| Mass | 3 | Sum, std deviation, mean |
| Pocket Propensity | 2 | [10] |
| Relative Accessible Surface Area | 1 | - |

#### 2.2 Environmental properties

Environmental descriptors are based on a series of radial functions that have previously been used to describe local atomic environments [11]. For a residue property $P$, environmental descriptors $D$ are computed according to,

$$D_i(r_c, \eta, \mu) = \sum_{\substack{j=1 \\ j \neq i}}^{N} P_j \cdot e^{-\eta(r_{ij}-\mu)^2} f_c(r_{ij}) \tag{3}$$

Here, $r_c$, $\eta$, and $\mu$ are hyperparameters and describe the distance and weighting across which the environment is considered. The pairwise distances between the center of masses of residue i and j are given by $r_{ij}$ and a cutoff function $f_c$ is computed as.

$$f_c(r_{ij}) = \begin{cases} \frac{1}{2}[\cos(r_{ij}\pi/r_c) + 1] & \text{if } r_{ij} \leq r_c \\ 0 & \text{if } r_{ij} > r_c \end{cases} \tag{4}$$

In addition to Equation 3, which represents an extrinsic summation, intrinsic environmental properties are also computed by normalizing the summation by the total property weights $W$

$$W_i = \sum_{\substack{j=1 \\ j \neq i}}^{N} e^{-\eta(r_{ij}-\mu)^2} f_c(r_{ij}) \tag{5}$$

**Table S3** Parameters used for environmental descriptors in NeCLAS. One asterisks describes the environmental descriptor parameters used in figure S4.

| $r_c$ | $\eta$ | $\mu$ | Type |
|---|---|---|---|
| 25 | 0.005 | 0 | sum |
| 18 | 0.05 | 0 | mean* |
| 18 | 1 | 7.5 | sum |
| 18 | 1 | 10 | sum |
| 18 | 1 | 12.5 | sum |

For each property, a total of five different environmental descriptors were computed by varying hyperparameters, as detailed in Table S3 and graphically shown in Figure S4.

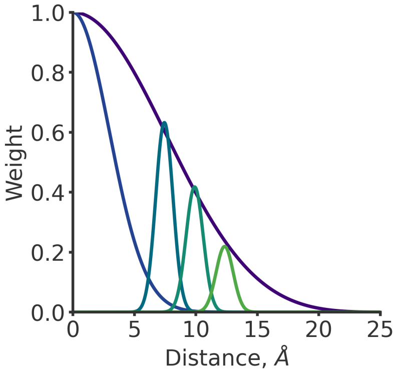

**Fig. S4** Visual representation of different weights used for environmental descriptors.

As discussed in the original work, no standard method exists for hyperparameter selection, but rather it is informed by knowledge of the system and obtaining a number of different coverage [11]. We choose 1.8 nm as the primary cutoff radius since it has been similarly employed with some success as a cutoff for Voronoi based environmental descriptors in similar protein-protein pairwise predictions [12] and nanoparticle machine learning studies [13]. Distances considered are from the center of masses, however, environmental interactions might occur between any atoms within two residues. Therefore, for a more complete description, we also consider at least one longer cutoff with an additional 0.7 nm which corresponds approximately to the difference in position between the center of mass and the outer heavy atoms in our datasets, for the most common CG size. Beyond that, we choose multiple values of $\mu$ to capture a number of different positional distances between the origin and the cutoff radius. To avoid biasing the data towards bound structures, environmental descriptors are only calculated using a single structure, not information of the bound structure.

These descriptors are also used to smooth our predictions. A number of other protein interaction prediction methods [12, 14] perform a basic smoothing of prediction results by considering nearby predictions. This has been empirically shown to improve results, as being surrounded by residues with a high probability of interaction is itself a good indicator of interaction [12]. To this end, we smooth our predictions by the weighted average prediction with weights determined by the environmental descriptors discussed above. We used,

$$P_i = (P'_i + D_i)/(W_i + 1) \tag{6}$$

where $P_i$ is the smoothed prediction, $D_i$ is given in equation 3, and $W_i$ is given in equation S4. The unsmoothed prediction $P'_i$ is given a maximum weight of 1. $r_c$, $\eta$, and $\mu$ are 25, 0.005, and 0 respectively.

### 3 Datasets

#### 3.1 Nanoparticle-Protein

The nanoparticle dataset used for testing is based on the atomistic details of organic nanoparticles provided by Costanzo *et al.* [15]. When identical cases of the same protein and nanoparticle interaction were provided, the case with more interactions was considered, and all possible sites were used for label generation. All examples that did not contain a valid nanoparticle structure were omitted. The unbound protein structures were identified by searching the RCSB database for proteins with matching sequences. Two structures (5ET3 and 5N10) contain no equivalent solved structure, and therefore the bound structure was used. Homology analysis between the training set and these nanoparticles is discussed in the main text.

**Table S4** List of protein-nanoparticle pairs used in protein-nanoparticle testing set. The PDB ID represents the bound complex. The unbound protein ID is the PDB ID of the unbound protein used. The NP ID is either the chain (if a single letter) or RCSB ligand ID of the nanoparticle. The number of sites used in coarse-graining the nanoparticle is also provided.

| PDB ID | Unbound Prot. | NP ID | Num. Sites |
|---|---|---|---|
| 3BCD | 3BCF | D | 6 |
| 3CYU | 1AVN | 1CR | 12 |
| 3EDK | 3EDD | C | 16 |
| 3TYI | 2MHM | T3Y | 8 |
| 3CZH | 3C6G | C | 7 |
| 4PRQ | 5WRB | T3Y | 8 |
| 5ET3 | 5ET3 | 60C | 6 |
| 5N10 | 5N10 | C8L | 16 |
| 5LFT | 2MHM | 6VB | 8 |
| 5LYC | 2MHM | 7AZ | 12 |
| 5KPF | 2MHM | 6VJ | 8 |
| 5OEH | 3IQU | 9SZ | 6 |
| 5MKA | 5MKB | B | 8 |
| 6HAH | 2MHV | FWQ | 12 |
| 6HAJ | 2MHV | EVB | 16 |
| 6HA4 | 2MHV | T3Y | 8 |
| 6EGY | 2MHM | B4T | 9 |
| 6RGI | 5T8W | FWQ | 12 |
| 6GL5 | 6F7Y | T3Y | 8 |
| 6GD6 | 2MHM | EVB | 16 |
| 6SUY | 2MHM | LVT | 8 |

### 4 Machine Learning

#### 4.1 Implementation Details

All NeCLAS training and validation data were randomly downsampled to a 3:1 ratio for negative and postive classes, respectively. No downsampling was performed on the testing datasets. All Neural Networks (NN) were implemented using TensorFlow/Keras (TF) 2.8.0 with eager execution disabled (PIPGCN was ported from TF1 to TF2). All NNs utilize variants of stochastic gradient descent, which have a probabilistic component that can provide slightly different results depending on the random seed used. Thus, we use the TensorFlow default of 32-bit floats for our 230-fold protein-protein cross validation. All NNs and XGBoost were trained using an RTX 3080 GPU and an Intel i9-11900KF CPU. During leave-one-protein-out (LOPO) cross validation, our permutation-invariant NN (NeCLAS) trained for approximately 5 hours at 40% GPU capacity, while XGBoost trained for approximately 13 hours at 100% GPU capacity.

For comparison, using PIPGCN's predefined split, PIPGCN utilized 75% GPU capacity with 10 minutes of training time, while NeCLAS utilized 40% of GPU capacity with 30 seconds of training time.

#### 4.2 Homology

We show the effect on our protein-nanoparticle evaluation results of removing proteins from our training set which share a SCOP [16, 17] family with any of the proteins in our protein-nanoparticle data set. The following complexes are removed: 1IB1, 1MZN, 1CMV, 1TMQ, 1BVN, 1KXQ, 1MLC, 2I25, 1DQJ, 1BVK, 1VFB, 2MTA. To account for different intial states in the neural network, we train 30 models without these homologs and discuss the results in the main text.

#### 4.3 NoPair

In the main text, we make the tacit assumption that partner-specific information from coarse-graining is necessary for improved prediction. Here, we use a simple ablation test to show that removing this additional information will significantly impact model performance. To generate a training dataset for this additional test, each protein in the DBD is considered separate from its partner when computing features, and each amino acid assigned a positive or negative label depending on whether it falls within a predefined interaction threshold of 6Å from any site on its partner molecule. The testing dataset is generated similarly, but we only consider the protein in each protein-nanoparticle complex. Removing the partner nanoparticle decreases the performance of NeCLAS with results discussed in the main text.

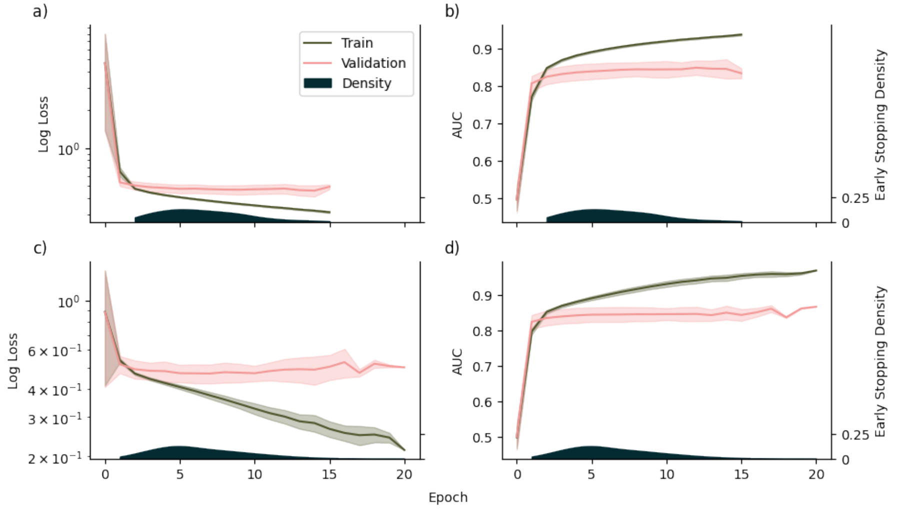

**Fig. S5** Subplots a) and b) show the loss and AUC curves, respectively, for training and validation during leave-one-out cross validation (LOO-CV). Subplots c) and d) show loss and AUC curves for the 10 models trained for predicting Nanoparticles. The variance of each metric is represented by the shaded region. Because we use early stopping, we include the distribution of training epochs reached before training was halted.

#### 4.4 Training

To ensure accurate and unbiased performance estimates for protein-protein interaction, we perform leave-one-protein-out cross validation over 230 proteins, where we iterate over each protein $P_i$ and remove all interactions involving $P_i$ from the training dataset. Our testing dataset is then constructed from all the interactions that include $P_i$. To provide robust estimates, we calibrate all preprocessing steps using only the training dataset, and we use a separate validation dataset of 16 proteins for early stopping. To construct the validation dataset, we partition the training dataset based on 3 "difficulty" levels (as defined by DBD database) and 3 "family" bins (enzyme, antibody, and other interactions), and excise 2 proteins from each difficulty and each family. This randomization reduces human bias when selecting validation sets, while ensuring that the validation set is diverse enough to facilitate the early stopping of our permutation invariant NNs. However, for results to be reproducible, the same random seed must be used every time the entire cross-validation cycle is run. XGBoost does not utilize validation data, so we trained XGBoost on both the training and validation datasets to provide a more robust performance estimate. If we did not train XGBoost on both the training and validation dataset, it would be at a disadvantage to the NN, which exploited the validation data for early stopping. All protein models were trained using Adam with a learning rate of $10^{-3}$ and a batch size of 256.

To show that our features are truly general, our nanoparticle interaction model uses only proteins for training and validation datasets. Therefore, to prevent overfitting, we significantly increased the batch size to $2^{15}$ and reduced the model size as seen in figure S6.

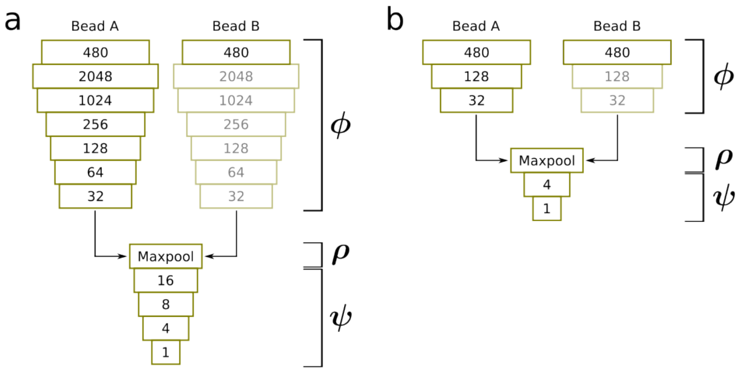

**Fig. S6** Architectural diagram of our permutation invariant NN. Each numbered layer represents a dense layer with that many weights, except for the first layer, which represents the input size. Transparent layers indicate shared weights. **a)** protein network with 2 input sites and 1 output task, **b)** miniaturized version of model a for nanoparticle prediction.

These modifications decrease training time, but also causes the model to occasionally get stuck in local minima. Similar to existing methods [18] (and unlike our protein-protein model), we rerun the same fit multiple times during training. Typically, these models are all used during inference, with their predictions being aggregated to form a single prediction. However, after training, we keep only the model with the highest (protein) validation AUC. Since the NeCLAS model is small, this provides high AUC scores for protein-nanoparticle interaction predictions, while incurring minimal overhead during training and no overhead during inference.

#### 4.5 Permutation Variance

Our permutation invariant neural network is inspired by the DeepSets [19] architecture. Given CG sites $A, B$ which yield descriptors $\mathbf{a}, \mathbf{b}$, we apply a permutation variant function $\boldsymbol{\phi}$ element-wise to the block vector $[\mathbf{a}\ \mathbf{b}]$ to produce a permutation-equivariant function $\hat{\boldsymbol{\phi}}([\mathbf{a}\ \mathbf{b}]) := [\boldsymbol{\phi}(\mathbf{a})\ \boldsymbol{\phi}(\mathbf{b})]$. We then apply a permutation invariant aggregator function $\boldsymbol{\rho}$ and a permutation variant function $\boldsymbol{\psi}$, resulting in a non-trivial permutation invariant function:

$$\boldsymbol{\sigma}([\mathbf{a}\ \mathbf{b}]) := \boldsymbol{\psi}(\boldsymbol{\rho}([\boldsymbol{\phi}(\mathbf{a})\ \boldsymbol{\phi}(\mathbf{b})]))$$

In our case, $\boldsymbol{\rho}$ is `Maxpool` and $\boldsymbol{\phi}, \boldsymbol{\psi}$ are Multilayer Perceptrons (MLP). We use ReLU activation for all layers except for the final layer of $\boldsymbol{\psi}$, which has sigmoid activation to enable binary predictions.

We demonstrate the practical implications of permutation variant models by comparing our permutation-invariant network against XGBoost models that are trained and evaluated on various CG sites orderings. We train one XGBoost model on the standard CG sites' ordering [**a b**], then evaluate it on both orderings ([**a b**] and [**b a**]). Training an XGBoost model on both orderings was projected to take 33 hours, so we sub-sampled this augmented training dataset to match the original training dataset size. Figures S7 and S8 show that XGBoost exhibits unstable predictions and a marked performance decrease when sites are permuted. Such permutations can even change the AUC of an individual complex by as much as 0.49. As a sanity check, we perform the same evaluation on our permutation-invariant neural network and find that the ordering has no effect on the model outputs.

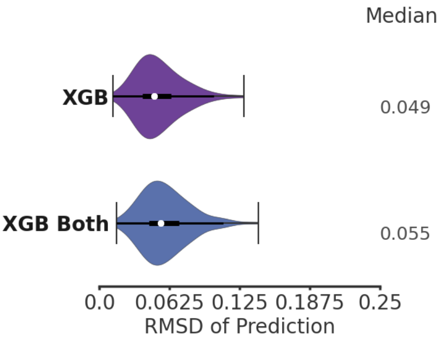

**Fig. S7** The root mean squared deviation (RMSD) between predictions over the range $[0, 1]$ for feature vectors [**a b**] and [**b a**]. We only show XGBoost trained on one direction and XGBoost trained on both directions, since the predictions of the permutation invariant network have a RMSD of 0

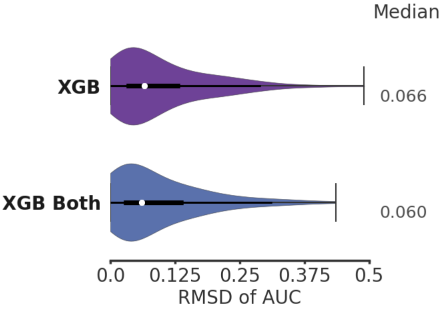

**Fig. S8** This figure shows the RMSD between the AUC of the prediction for [**a b**] and the AUC of the prediction for [**b a**]. The permutation-invariant network is not included, as it achieved 0 RMSD of AUC for each complex.

#### 4.6 Other Methods

Comparisons with competing methods were carried out using the published implementations of the respective method using as inputs the unbound protein structures except for Unified [18], which uses the protein and its partner (*e.g.*, nanoparticle). A standard evaluation at the amino acid level was carried out for all methods, however, not all predictions were provided in this format. Therefore, for Fpocket [20], we applied a validation approach used previously, which converts atom predictions to residue predictions by assigning the maximum value of its constituent atoms [21]. For COACH [22], the organic interaction prediction was used as the interaction probability, since we consider entirely organic systems. Both Unified and NeCLAS have data preprocessing steps. NeCLAS's preprocessing step requires approximately 2 hours on a consumer CPU. However, due to its long computation time, Unified's descriptors were computed on the Michigan ARC computing cluster with 4-16 cores depending on the protein size. Although proteins were processed in parallel, computation times averaged from 30 minutes to several hours, each. 1N2C in particular required approximately 2 days of computing time. Unified was unable to preprocess 2 proteins (1OPH, 1F51), so we left out these proteins during evaluation.

### 5 Test Cases. Additional data

#### 5.1 PSMα1 and GQD Interactions

Figure S9 provides a detailed depiction of the coarse-graining of the GQD. There are three internal sites and 9 edge sites, each of which are nearly identical to each other.

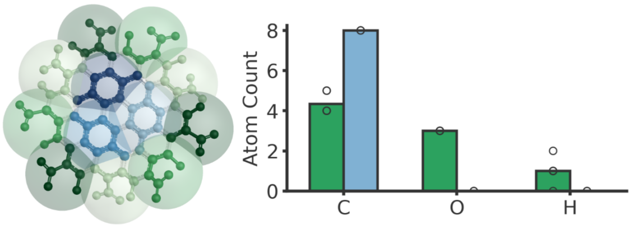

**Fig. S9 Coarse-graining of GQD**. Left: Depiction of coarse-graining used in GQD-PSMα1 study. Right: Atomic composition of GQD CG sites. The bar represents mean atom count and circles individual values. Green represents exterior CG sites, blue represents interior.

We also tested the correlation between NeCLAS's predictions and different definitions of contact time (Fig. S10).

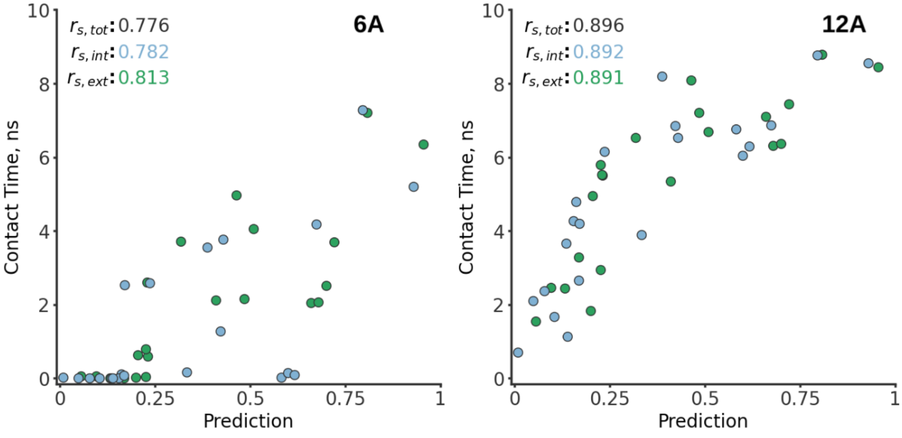

**Fig. S10** Contact time vs. interaction prediction for two different thresholds of contact time, denoted in the top right of each panel (6 Å and 12 Å; main text shows 10 Å). Edge sites are green and center sites are blue. Spearman correlation is given in the top left.

We define interaction by the contact time, which is the total simulation time when an amino acid residue and GQD CG sites are within a distance threshold. Of note, due to the symmetrical nature of GQD, the molecular dynamics simulation results showed a high level of interaction between some sites and a low interaction between other nearly identical ones, depending on the configuration of the protein and nanoparticle. Therefore, when comparing the contact time for each protein residue of PSM$\alpha$1, we consider the internal and edge sites, which spend the most time within the distance threshold. While in the main text, we use a threshold of 1 nm, as it is the shortest distance that produces non-zero contact time for all sites, results for 0.6 nm and 1.2 nm demonstrate that the correlation between prediction and contact time are still highly correlated.

#### 5.2 GQD Aggregation

Molecular dynamics simulations were performed with either Large-scale Atomic/Molecular Massively Parallel Simulator (CG simulations, software version 29 Sep 2021 - Update 2) [23] or Nanoscale Molecular Dynamics Program (all-atom simulations, software version 2.13) [24].

For intramolecular interactions in the CG simulations we used harmonic potentials for bonds, angles, dihedral, and improper, using all atom equilibrium distance/angles as equilibrium values and constants of 150/75 kcal/mol/Å$^2$ for bonds, 100/50 kcal/mol for angles (in both cases the first value is for rigid aromatic atoms, the second for everything else), and 70/35/17.5 kcal/mol for dihedral and improper, based on the amount of atoms that were part of a rigid aromatic subgroup.

For intermolecular interactions, we used

$$E(r) = 4\epsilon\sqrt{p}\left\{\left[\frac{(1-p)^2}{2} + \left(\frac{r}{\sigma}\right)^6\right]^{-2} - \left[\frac{(1-p)^2}{2} + \left(\frac{r}{\sigma}\right)^6\right]^{-1}\right\} \quad (7)$$

where $p$ ($\in [0,1]$) is the prediction value from NeCLAS. $\sigma$ (nm) and $\epsilon$ (kcal/-mol) were kept the same for all the CG sites of a given GQD. $\sigma$ was chosen to be 4 nm based on the distances observed in [25]. $\epsilon$ was estimated by matching the minimum for the potential in Eq. 7 for benzene-benzene interactions (NeCLAS prediction $\approx 0.2$) with the energy value of the potential for the closest minimum for the interactions between two CG benzene molecules, $\epsilon_{benzene} = 0.5$ kcal/mol (from [26]). The match produces a value of $\epsilon \approx 1.11$ kcal/mol. These two values were used for all the intermolecular potential, while $p$ was obtain from NeCLAS's predictions.

For all atom simulations, we run the fully solvated GQDs in canonical ensemble, starting from the final conformation produced in our previous work [25]. Complete details about the protocol and force field can be found there.

### References

[1] Martinetz, T., Berkovich, S. & Schulten, K. 'Neural-gas' network for vector quantization and its application to time-series prediction. *IEEE Transactions on Neural Networks* **4** (4), 558–569 (1993) .

[2] Arthur, D. & Vassilvitskii, S. *K-means++: The advantages of careful seeding*, SODA '07, 1027–1035 (Society for Industrial and Applied Mathematics, USA, 2007).

[3] Arkhipov, A., Freddolino, P. L. & Schulten, K. Stability and Dynamics of Virus Capsids Described by Coarse-Grained Modeling. *Structure* **14** (12), 1767–1777 (2006) .

[4] Dolnicar, S. & Leisch, F. Evaluation of structure and reproducibility of cluster solutions using the bootstrap. *Marketing Letters* **21**, 83–101 (2010) .

[5] Stanton, D. T. & Jurs, P. C. Development and use of charged partial surface area structural descriptors in computer-assisted quantitative structure-property relationship studies. *Analytical Chemistry* **62** (21), 2323–2329 (1990) .

[6] Stanton, D. T., Egolf, L. M., Jurs, P. C. & Hicks, M. G. Computer-assisted prediction of normal boiling points of pyrans and pyrroles. *Journal of Chemical Information and Computer Sciences* **32** (4), 306–316 (1992) .

[7] Todeschini, R. & Gramatica, P. The Whim Theory: New 3D Molecular Descriptors for Qsar in Environmental Modelling. *SAR and QSAR in Environmental Research* **7** (1-4), 89–115 (1997) .

[8] Sanner, M. F., Olson, A. J. & Spehner, J.-C. Reduced surface: An efficient way to compute molecular surfaces. *Biopolymers* **38** (3), 305–320 (1996) .

[9] Pintar, A., Carugo, O. & Pongor, S. CX, an algorithm that identifies protruding atoms in proteins. *Bioinformatics* **18** (7), 980–984 (2002) .

[10] Kawabata, T. Detection of multiscale pockets on protein surfaces using mathematical morphology. *Proteins: Structure, Function, and Bioinformatics* **78** (5), 1195–1211 (2010) .

[11] Gastegger, M., Schwiedrzik, L., Bittermann, M., Berzsenyi, F. & Marquetanda), P. wacsf—weighted atom-centered symmetry functions as descriptors in machine learning potentials. *J. Chem. Phys.* **148** (2018) .

[12] ul Amir Afsar Minhas, F., Geiss, B. J. & Ben-Hur, A. Pairpred: Partner-specific prediction of interacting residues from sequence and structure.

*Proteins* **82**, 1142–1155 (2014) .

[13] Yan, X. *et al.* *In silico* profiling nanoparticles. *Nanoscale* **11** (17), 8352–8362 (2019) .

[14] Sanchez-Garcia, R., Sorzano, C., Carazo, J. M. & Segura, J. Bipspi: a method for the prediction of partner-specific protein-protein interfaces. *Bioinformatics* **35** (3), 470–477 (2019) .

[15] Costanzo, L. D. & Geremia, S. Atomic details of carbon-based nanomolecules interacting with proteins. *Molecules* **25** (15), 3555 (2020) .

[16] Andreeva, A., Howorth, D., Chothia, C., Kulesha, E. & Murzin, A. G. SCOP2 prototype: a new approach to protein structure mining. *Nucleic Acids Research* **42** (D1), D310–D314 (2013) .

[17] Andreeva, A., Kulesha, E., Gough, J. & Murzin, A. G. The SCOP database in 2020: expanded classification of representative family and superfamily domains of known protein structures. *Nucleic Acids Research* **48** (D1), D376–D382 (2019) .

[18] Cha, M. *et al.* Unifying structural descriptors for biological and bioinspired nanoscale complexes. *Nature Computational Science* **2**, 243–252 (2022) .

[19] Zaheer, M. *et al.* Guyon, I. *et al.* (eds) *Deep sets.* (eds Guyon, I. *et al.*) *Advances in Neural Information Processing Systems*, Vol. 30 (Curran Associates, Inc., 2017).

[20] Le Guilloux, V., Schmidtke, P. & Tuffery, P. Fpocket: An open source platform for ligand pocket detection. *BMC Bioinformatics* **10** (1), 168 (2009) .

[21] Ezzat, A. & Kwoh, C. K. Comparison of Structure-based Tools for the Prediction of Ligand Binding Site Residues in Apo-structures. *Procedia Computer Science* **11**, 115–126 (2012) .

[22] Yang, J., Roy, A. & Zhang, Y. Protein-ligand binding site recognition using complementary binding-specific substructure comparison and sequence profile alignment. *Bioinformatics (Oxford, England)* **29** (20), 2588–2595 (2013) .

[23] Plimpton, S. Fast parallel algorithms for short-range molecular dynamics. *Journal of Computational Physics* **117** (1), 1–19 (1995) .

[24] Phillips, J. C. *et al.* Scalable molecular dynamics with NAMD. *Journal of Computational Chemistry* **26** (16), 1781–1802 (2005) .

[25] Elvati, P., Baumeister, E. & Violi, A. Graphene quantum dots: effect of size, composition and curvature on their assembly. *RSC Advances* **29** (2017) .

[26] Elvati, P. *Computer Simulations of Fuel Cells: Modeling of Surfactants and Polymeric Membranes for Fuel Cells Applications* (LAP LAMBERT Academic Publishing, Saarbrücken, Germany, 2012).
